# Distribution and association of biophysical factors with onion (*Allium cepa* L.) downy mildew (*Peronospora destructor*) disease epidemics in northwestern Ethiopia

**DOI:** 10.1101/2025.08.26.672276

**Authors:** Mitku Bitew, Addisu Mandefro, Yehizbalem Azmeraw

## Abstract

Onion (*Allium cepa* L.) is a significant food bulb crop that can be used for medicine, diverse recipes, and income. However, downy mildew disease limits productivity and production of onion in onion growing-regions. Thus, this survey was conducted to assess the distribution and intensity of onion downy mildew and determine the associations of disease parameters with biophysical factors. During the 2024/2025 onion cropping season, 130 onion fields were surveyed in order to measure biophysical and disease data. The associations of disease parameters and biophysical factors were analyzed using the binary logistic regression model by employing the SAS GENEMODE procedure. The survey’s findings verified that downy mildew was 100% prevalent. The highest disease intensity was assessed from Fogera (50.68% and 23.48%) and Libokemkem (45.16% and 21.62%) districts, respectively. High disease incidence (>40%) and severity (>20%) were strongly associated with maturity crop growth stages, field size (>0.25 ha), blub previous crop, late October transplanting, less than four times fungicide application and less than three times land preparation. Lower disease incidence (≤ 40%) and severity (≤ 20%) had a strong association with early December transplanting, more than three times land preparation, fungicide spray (>4 times), field size (≤ 0.25 ha), fields previously planted with cereals, could be considered as relative management options to minimize the disease’s effects in onion growing areas of northwest Ethiopia and other similar onion cultivating areas. In addition, In order to develop effective management strategies, future research may examine pathogen variability.

## 1. INTRODUCTION

The onion (*Allium cepa* L.), a bulb crop in the Amaryllidaceae family and essential spices of vegetable crops grown in many countries of the world. It originated in southwest Asia and is the most important commercial bulb crop in the vegetable bulb crop group. In terms of yearly global production of vegetable crops, onion ranks second after tomato [1]. Most smallholder farmers and a few commercial producers cultivate it year-round for both domestic and export markets, mostly under irrigation and to a lesser degree under rain-fed conditions [2, 3].

According to [1] report, onion production exceeds 93.22 megatons (Mt) in world annually. China is the world’s largest producer of onions with production of 23.91 Mt, followed by India (19.42 Mt). In Ethiopia, about 3.46 megatons (Mt) were produced in 38,952.58 ha land in 2020/21 cropping season of with an average yield of 8.8t ha^-1^ [4], which is lower than the other onion-producing countries. The onion is a crop that thrives in semi-arid regions, high elevations, and tropical climates. Climate characteristics that are favorable for onion production in Ethiopia include temperatures between 17 and 27 °C, 500 to 700 mm of annual rainfall, and elevations up to 2200 m. a. s. l. Onion production can also be achieved in fertile sandy and silt loam soils with a pH range of 6.0 to 6.8, while excessive rainfall and extremely hot or cold temperatures can reduce yield [5].

Onion is one of the most significant vegetable crops grown globally for food, as well as a relative and a valuable addition to many recipes [6, 7, 8]. In world including Ethiopia, The crop is valued for its distinct pungency and form essential ingredients for flavoring varieties of dishes, sauces, soup, sandwiches and snacks as onion rings and also exhibits a number of therapeutic properties, such as antibacterial, antifungal, anti-helminthic, anti-inflammatory, antiseptic, antispasmodic, etc. and it contributes to the national economy through export products like, bulbs and cut flowers to different countries of the world [9, 10, 11, 12]. Because of these significant advantages, onion production is growing in various agro-ecologies across the nation in small-scale production systems as a part of commercialization.

Despite its importance for household consumption, medicinal uses, income sources and contribution to the national economy through exports, the onion production and productivity are constrained by several abiotic and biotic stresses [13]. The national average production of onion under farmers’ situations is considerably deficit in Ethiopia (8.9 t per ha^-1^) [4], which are significantly lower than global average (19.1t ha^−1^). The Amhara region is a major contributor, accounting for about 50% of the national production with a yield of 12.3 tons per hectare [14]. This low productivity is mainly associated with different diseases and insect pests. In this regard, diseases, such as purple blotch (*Alternaria porri*) and downy mildew (*Peronospora destructor* are the major economically important diseases that lowering onion yields [15, 16, 17, 18, 19].

Among diseases, downy mildew is one of the most widespread and devastating disease and cause highest yield losses in onion producing areas in the world [15, 20, 21, 22]. It’s characterized by yellowing and dieback of leaves, with a white and then purple fungal growth on the leaf surface. It can also cause premature sprouting or shriveling of bulbs. It is a water mold or oomycete disease of alliums caused by *Peronospora destructor*. It causes irregular foliar lesions that begin as pale-green, and then progress to yellow or brown necrotic tissue. Eventually, lesions coalesce and lead to the collapse of the leaf. During periods of high moisture, fuzzy, gray-to-violet sporangia appear on leaf surfaces. These symptoms can also be seen on seed stalks and flowers [23, 24]. When conditions are favorable for the disease to occur, downy mildew outbreaks can cause yield decreases of onion bulbs of 30% to 75% [25, 26]. In northwestern Ethiopia, yield loss due to downy mildew has recorded up to 80% of the total production with the relative occurrence of the disease [27].

Various management strategies are employed to lessen the disease’s transmission and infection sources. The management of onion downy mildew involves a number of strategies, including the Planting timing, resistant cultivars [28, 29]. Soil drainage [30], seed or bulb health stock [31], wind block and the elimination of infected plants [32, 33]. However, the most effective means of controlling the downy mildew is the use of fungicides [34].

Despite the fact that the disease is a significant barrier to onion production, intensive field survey on the disease epidemic is not done in the study areas rather than preliminary survey and fungicide evaluation studies. Thus, compressive field survey on downy mildew epidemics in onion producing-areas in northwestern Ethiopia could aid to provide information on the association of disease epidemic with cultural and environmental factors and to identify the existing management options at farmers’ level. Therefore, the study was conducted to assess the distribution and intensity of onion downy mildew and determine the association of the disease parameters with biophysical factors in northwestern Ethiopia.

## 2. MATERIALS AND METHODS

### 2.1. Description of the study areas

A field survey was conducted in the South and Central Gondar Zones of the Amhara Region Ethiopia, which are major onion-producing regions. During the 2024–2025 onion growing season, field assessments were conducted in three South Gondar districts (Fogera, Libokemkem, and Farta) and one Central Gondar zone district (Gondar Zuria), as shown in Figure 1. The districts are situated between latitudes 8°45’ and 13°45’ N and longitudes 36° 20’ and 40° 20’ E in the northwest of the Amhara Region. The districts were found between elevations of 1790 and 2186 meters above sea level, and they receive 1000 to 1600 millimeters of rainfall annually [35]. The study areas’ have an average temperature of ranged from 17.8 to 27.05 °C. The two main soil types found in the survey areas are nitisol and vertisol. Rainfall and minimum and maximum temperature of the districts were obtained from meteorological stations and are presented in Figure 2.

**Figure 1.**
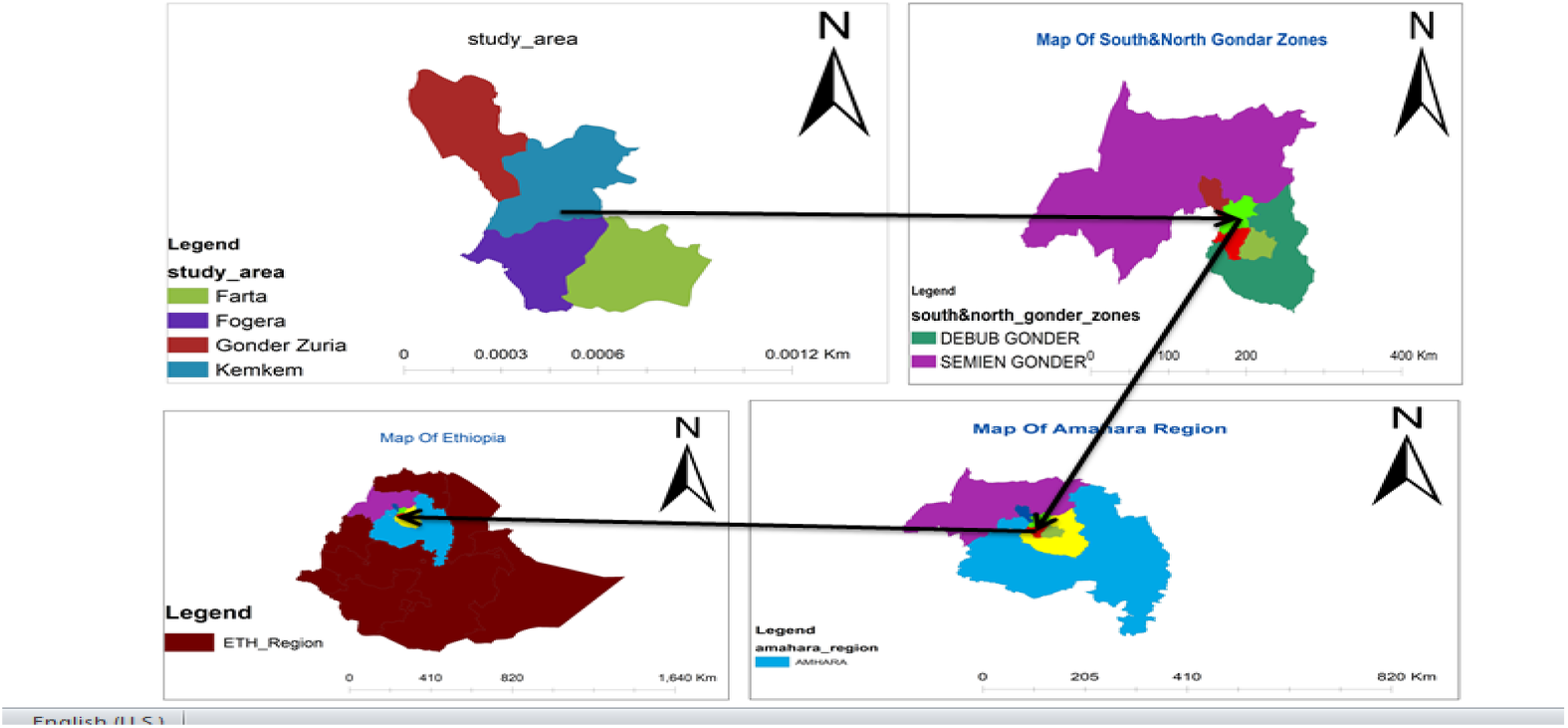
Map showing the districts for onion downy mildew diseases study in northwestern Ethiopia in 2024/25 cropping season.

**Figure 2.**
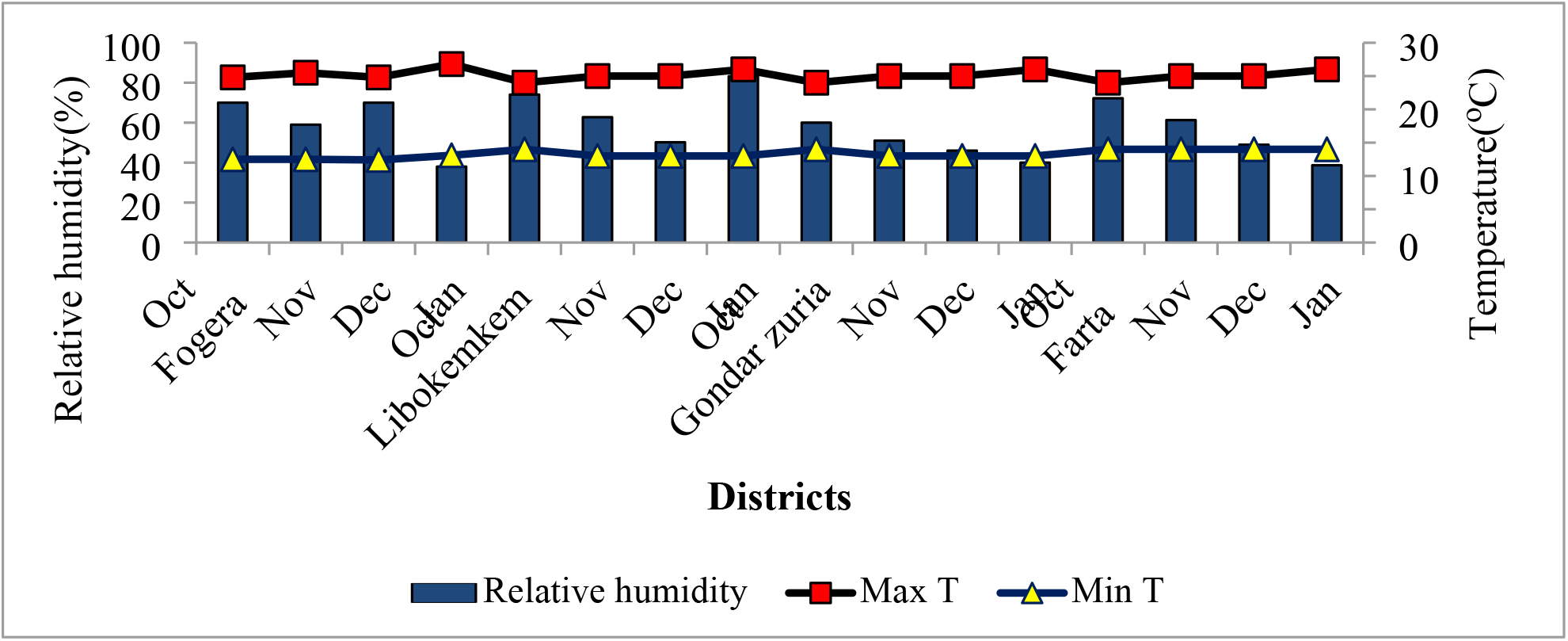
Average Temperatures (°C) and Relative humidity (%) of surveyed districts in northwestern of Ethiopia from sowing to surveyed periods of 2024/25 onion growing season.

### 2.2. Sampling procedure and diseases assessment

The districts were specifically selected on the basis of their potential for onion production. Five to seven representative Farmers Associations were selected from each district, and each Farmers’ Association had six to ten fields examined. Random sampling was used to select onion fields at 4-to 5-kilometer intervals. The plant population was assessed using a 1 m^2^ quadrat arranged in a “X” layout with 5 m between each field. The mean plant population densities were calculated by averaging the number of populations in the quadrat for analysis. Five spots were evaluated diagonally in each field (using a 1m2 quadrat) to avoid bias in recording the disease’s severity. A total of 130 fields were assessed across four districts throughout the survey.

The number of onion plants in each quadrat was counted, and the number of infected plants was noted. Ten randomly selected plants were evaluated for disease severity in each quadrat using a 1–9 rating system [34]; where, 1 = no disease symptoms, 2 = only a few leaves 1% affected, 3 = less than half of the plants (5%) affected, 4 = most of the plants affected, 10% attack is restricted to one leaf per plant, 5 = all plants attack, (20%) attack restricted to one or two leaves, 6 = three to four leaves of each 50% of plants affected, crops looks fairly green, 7 = all leaves affected 75% of crop gives blight appearance, 8 = all leaves severely affected 90%, greenness restricted to central shoot only, 9 = foliage completely blighted 100%. Then, the severity scale was converted to percent severity index (PSI) [36]. Disease prevalence [37], incidence [38], and percent severity index in each sampled field were calculated as follows:

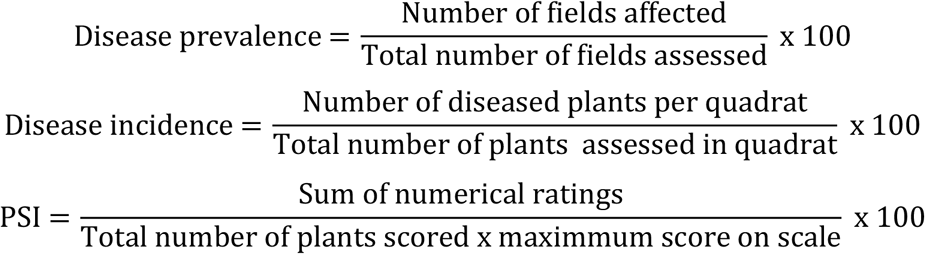

Additional data, including previous field history, seed source, transplanting date, land preparation, and frequency of fungicide application, were gathered during the survey by interviewing farmers using questionnaires. Visual observation was used to record the field size and crop growth stage. The Global Positioning System (GPS) was used to record the altitude, latitude, and longitude.

### 2.3. Data analysis

Descriptive analysis was used to show the distribution and association of the incidence and severity of onion downy mildew disease and biophysical parameters in each district. To develop a binary dependent variable, class boundaries were established for both disease incidence and severity into two separate groups’ binomial qualitative data: ≤ 40 and > 40% were chosen for disease incidence, while ≤ 20 and >20% were selected for severity. To represent the bivariate distribution of fields based on data classifications, a contingency table of disease incidence and severity as well as the independent variables was constructed (Table 1).

**Table 1.** Categorization of variables used in analysis for the distribution of onion downy mildew epidemics in four districts (*n* = 130) of northwestern Ethiopia, during the 2024/25growing season.

| Variable | Variable class | No. of fields | Incidence (%) |  | Severity (%) |  |
| --- | --- | --- | --- | --- | --- | --- |
|  |  |  | ≤ 40 | > 40 | ≤ 20 | > 20 |
| District | Fogera | 62 | 23 | 39 | 28 | 34 |
|  | Libokemkem | 30 | 8 | 22 | 11 | 19 |
|  | Gondar Zuria | 26 | 13 | 13 | 12 | 14 |
|  | Farta | 12 | 4 | 8 | 7 | 5 |
| Altitude (m. a. s. l.) | ≤ 1800 | 81 | 32 | 49 | 36 | 45 |
|  | > 1800 | 49 | 20 | 29 | 22 | 27 |
| **Seed source** | Farmer saved | 60 | 24 | 46 | 32 | 38 |
|  | Local market | 70 | 24 | 36 | 26 | 34 |
| **Plant population<sup>a</sup>** | Low | 32 | 14 | 18 | 17 | 15 |
|  | High | 98 | 34 | 64 | 41 | 57 |
| **Growth stage** | Vegetative | 56 | 27 | 29 | 31 | 25 |
|  | Bulbing | 42 | 15 | 27 | 15 | 27 |
|  | Maturity | 32 | 10 | 22 | 12 | 20 |
| **Land preparation** | ≤ 3 times | 58 | 24 | 34 | 29 | 29 |
|  | > 3 times | 72 | 24 | 48 | 29 | 43 |
| **Field size (ha)** | ≤ 0.25 | 51 | 20 | 31 | 30 | 21 |
|  | > 0.25 | 79 | 28 | 51 | 28 | 51 |
| **Transplanting date<sup>b</sup>** | Late October | 62 | 22 | 40 | 25 | 37 |
|  | Early December | 78 | 26 | 42 | 33 | 35 |
| **Previous crop** | Legume & Cereal | 49 | 24 | 25 | 23 | 26 |
|  | Bulb | 61 | 24 | 57 | 35 | 46 |
| **Fungicide application<sup>c</sup>** | ≤ 4 times | 56 | 21 | 35 | 29 | 27 |
|  | >4 times | 74 | 27 | 47 | 29 | 45 |

<sup>a</sup>Fields with plant population < 55 and ≥ 55 onion plants per 1m<sup>2</sup> quadrat were considered as low and high populated, respectively. <sup>b</sup>Late October planting starts from October 15 to 30 and planting of seedling made in the beginning of December was considered as early December planting. <sup>c</sup>Fungicide applications refer to spray frequencies from once to fourth consider as ≤ 4 times and above 4 times sprays as > 4 times.

To investigate the associations between biophysical factors and onion downy mildew disease, a binary logistic regression model was applied [39], using SAS procedure of GENMOD (SAS, 2016). The importance of the independent variables was evaluated twice in terms of their effect on the incidence and severity. First, using a single-variable model, the relationship between each independent variable and the incidence and severity of downy mildew was analyzed. Second, when all other factors in the model were entered first and last, the relationship between an independent variable and the incidence or severity of the disease was investigated. Finally, a reduced multiple-variable model was constructed by adding those independent variables that had a significant association with the incidence or severity of the disease. The odds ratio, which is understood as the relative risk, was calculated by exponentiating the parameter estimates for comparing the effect based on a reference point. A complete analysis of the deviance reduction was calculated for each variable as it was added to the reduced multiple-variable model [39]. The GENMOD process uses a maximum likelihood estimation of the parameter to fit a generalized linear model to the data. The comparison between the single and multiple variable models was done using the deviation, which is the logarithm of the ratio of two likelihoods. To assess the significance of the variable, the difference between the likelihood ratio tests (LRTs) was compared to the Chi-square value [40].

## 3. RESULTS

### 3.1. General descriptions of surveyed areas

During the survey, all of the assessed fields were free from weeds, and located in the altitude range of 1790 to 2186 m. a. s. l. The crops commonly grown in the districts were rice, potatoes, sorghum, chickpeas, *teff*, wheat, barley, garlic and chickpeas. Every assessed field were sprayed with fungicide, and planted in rows with varied source of seeds. About 46.15% fields were planted with farmer saved seeds and the remaining 53.85% fields were covered by seeds bought from local market. Depending on the farmer, different fields have different land preparation procedures before planting onions and 55.38% of the fields were plowed more than 3 times. Mainly, onion planting took place from October to December. Different growth (vegetative, bulbing and maturity) stages were encountered during the survey periods, which accounted for 43.07, 32.31 and 24.62% of these growth stages, respectively (Table 1).

### 3.2. Prevalence, incidence and severity of onion downy mildew

The onion downy mildew disease spread in various ways throughout different regions. The disease prevalence was recorded in all surveyed districts. The highest (50.68%) mean disease incidence was observed in Fogera district, followed by Libokemkem (45.16%) and the lowest (35.65%) disease incidence was recorded in Farta district. Similarly, higher disease severity was assessed from Fogera and Libokemkem districts (Table 3). Higher Onion downy mildew disease incidence (44.25%) and severity (22.27%) were assessed at fields located at lower altitudes (≤1800 m. a. s. l.) than higher (>1800 m. a. s. l.) altitudes (Table 2). Regarding seed source, higher downy mildew incidence (42.37%) and severity (23.67%) were noted from fields planted with seeds bought from market than those fields planted with farmer saved seeds. Variability in onion downy mildew intensity was observed among onion growth stages. In this regard, onion fields at maturity growth stage resulted in higher disease incidence (46.91%) and severity (26.97%) than bulbing and vegetative growth stages (Table 2) Land preparation (≤ 3 times) relatively increased onion downy mildew incidence from 39.16 to 41.08% and severity from 11.04 to 12.47 compared to land ploughed more three times.

**Table 2.** Incidence and severity (mean ± SE) of onion downy mildew for different independent variables in northwestern Ethiopia, during the 2024/25 growing season.

| Variable | Variable class | Incidence (%) | SE | Severity (%) | SE |
| --- | --- | --- | --- | --- | --- |
| **District** | Fogera | 50.68 | 15.23 | 23.48 | 10.73 |
|  | Libokemkem | 45.16 | 19.43 | 21.62 | 14.26 |
|  | Gondar Zuria | 37.65 | 16.59 | 21.38 | 7.53 |
|  | Farta | 35.86 | 12.56 | 21.23 | 10.23 |
| **Altitude (m. a. s. l.)** | $\leq 1800$ | 44.25 | 15.89 | 22.27 | 8.82 |
| | $> 1800$ | 37.79 | 18.08 | 21.78 | 13.53 |
| **Seed source** | Farmer saved | 38.39 | 16.98 | 20.51 | 13.03 |
|  | Local market | 42.37 | 16.85 | 23.67 | 10.37 |
| **Plant population<sup>a</sup>** | Low | 37.10 | 15.82 | 20.19 | 10.57 |
|  | High | 41.25 | 17.29 | 22.55 | 12.36 |
| **Growth stage** | Vegetative | 35.97 | 18.31 | 17.72 | 10.57 |
|  | Bulbing | 40.81 | 11.92 | 23.81 | 10.94 |
|  | Maturity | 46.91 | 18.34 | 26.97 | 13.18 |
| **Land preparation** | $\leq 3$ times | 41.08 | 17.21 | 23.57 | 12.47 |
| | $> 3$ times | 39.16 | 16.75 | 19.99 | 11.04 |
| **Field size (ha)** | $\leq 0.25$ | 37.69 | 14.15 | 19.42 | 10.89 |
| | $> 0.25$ | 41.86 | 18.46 | 23.61 | 12.36 |
| **Transplanting date<sup>b</sup>** | Late October | 41.17 | 19.66 | 20.69 | 12.74 |
|  | Early December | 39.36 | 13.52 | 23.37 | 10.92 |
| **Previous crop** | Legume & Cereal | 36.68 | 14.63 | 22.56 | 13.21 |
|  | Bulb | 42.37 | 17.99 | 21.61 | 11.18 |
| **Fungicide application<sup>c</sup>** | $\leq 4$ times | 40.73 | 11.52 | 24.49 | 12.04 |
| | $> 4$ times | 39.57 | 22.34 | 18.65 | 11.07 |

<sup>a</sup>Fields with plant population $< 55$ and $\geq 55$ plants per 1m<sup>2</sup> quadrat were considered as low and high populated. <sup>b</sup>Late October planting starts from October 15 to 30 and early December planting refers planting of seedlings in the beginning of December. <sup>c</sup>Fungicide applications refer to spray frequencies from once to fourth considered as $\leq 4$ times and above 4 times sprays as $> 4$ times. SE = Standard error.

**Table 3.** Logistic regression model for onion downy mildew incidence and severity, and likelihood ratio test on independent variables in northwestern Ethiopia, during the 2024/25 onion growing season.

| Independent Variable | df | Onion downy mildew incidence, LRT<sup>a</sup> |  |  |  | Onion downy mildew PSI, LRT |  |  |  |
| --- | --- | --- | --- | --- | --- | --- | --- | --- | --- |
|  |  | Type 1 analysis (VEF) |  | Type 3 analysis (VEL) |  | Type 1 analysis (VEF) |  | Type 3 analysis (VEL) |  |
| | | DR | Pr> $\chi^2$ | DR | Pr> $\chi^2$ | DR | Pr> $\chi^2$ | DR | Pr> $\chi^2$ |
| **District** | 3 | 5.88 | 0.1175 | 5.85 | 0.1193 | 0.52 | 0.9139 | 4.41 | 0.2203 |
| **Growth stage** | 2 | 41.47 | <.0001 | 18.57 | <.0001 | 99.25 | <.0001 | 20.96 | <.0001 |
| **Land preparation** | 1 | 0.16 | 0.0004 | 0.05 | 0.0007 | 84.88 | <.0001 | 15.73 | <.0001 |
| **Field size (ha)** | 1 | 0.63 | 0.4264 | 0.63 | 0.426 | 17.98 | <.0001 | 8.07 | 0.0001 |
| **Seed source** | 1 | 8.89 | 0.0029 | 9.02 | 0.0027 | 132.09 | <.0001 | 29.21 | <.0001 |
| **Previous crop** | 1 | 9.55 | 0.0020 | 9.55 | <.0001 | 37.80 | <.0001 | 14.65 | 0.0001 |
| **Transplanting date** | 1 | 9.37 | 0.0022 | 1.95 | 0.1624 | 15.10 | 0.0001 | 11.43 | 0.0007 |
| **Altitude** | 1 | 0.08 | 0.7736 | 0.01 | 0.9046 | 1.84 | 0.1754 | 1.84 | 0.1754 |
| **Fungicide application** | 1 | 14.04 | 0.0002 | 16.78 | 0.0020 | 5.35 | 0.0207 | 9.57 | 0.0020 |
| **Plant population** | 1 | 7.34 | 0.0067 | 0.37 | 0.3230 | 35.55 | <.0001 | 36.22 | <.0001 |

<sup>a</sup>LRT = Likelihood ratio test. VEF = Variable entered first in the model. VEL = Variable entered last in the model. DR = Deviance reduction. Pr = Probability of a $\chi^2$ value exceeding the deviance reduction. $\chi^2$ = Chi-square. df = Degrees of freedom and PSI = Percent severity index.

Variable disease intensity was recorded in terms of onion transplanting dates, where late October planting was relatively suitable for downy mildew disease epidemic with 41.17% mean incidence and 12.74% severity. Concerning previous crop, the highest incidence (42.37%) and severity (13.21%) with large lesion symptoms were observed on onion growing previously after blubs than cereals and legumes (Table 3). Fields with fungicide spray (≤4 times) had higher disease incidence and severity than onion fields with spray (> 4 times) (Table 3).

### 3.3. Association of onion downy mildew incidence and severity with biophysical factors

The associations of all biophysical factors with disease incidence and severity are described in Table 4. The biophysical factors, such as growth stage, land preparation, seed source, previous crop history and fungicide application were significantly (P< 0.05) associated with onion downy mildew disease intensity when entered first and last into the logistic regression model and in addition those associated with intensity, field size, transplanting date and plant population attained its significant association with onion downy mildew severity when entered first and last into logistic regression model (Table 3).

**Table 4.** Analysis of deviance, natural logarithms of odd ratio and standard error of onion downy mildew incidence and likelihood ratio test on independent variables in northwestern Ethiopia, during the 2024/25 growing season.

| Variable<sup>a</sup> | Residual Deviance<sup>b</sup> | Downy mildew incidence, LRT<sup>c</sup> |  | Variable class | Estimate<sup>d</sup> | SE | Odds ratio |
| --- | --- | --- | --- | --- | --- | --- | --- |
| | | DR | Pr> $\chi^2$ | | | | |
| **Intercept** | 174.1400 |  |  | Intercept | 1.7807 | 0.81 | 5.93 |
| **District** | 168.2586 | 1.90 | 0.11 | Fogera | 1.1787 | 1.16 | 3.25 |
|  |  |  |  | Libokemkem | 1.0279 | 0.83 | 2.79 |
|  |  |  |  | Gondar Zuria | 0.7518 | 0.72 | 2.12 |
|  |  |  |  | Farta | 0* |  | 1 |
| **Growth stage** | 125.2779 | 21.69 | <.0001 | Vegetative | -4.0792 | 0.99 | 0.016 |
|  |  |  |  | Bulbing | -2.6182 | 0.78 | 0.072 |
|  |  |  |  | Maturity | 0* |  | 1 |
| **Land preparation** | 152.2489 | 11.51 | 0.0004 | ≤ 3 times | 1.8400 | 0.59 | 6.29 |
|  |  |  |  | > 3 times | 0* |  | 1 |
| **Field size** | 123.4532 | 0.58 | 0.44 | ≤ 0.25 | -0.4769 | 0.64 | 0.62 |
|  |  |  |  | > 0.25 | 0* |  | 1 |
| **Previous crop** | 152.1389 | 0.01 | 0.0022 | Legume&cereal | -0.0635 | 0.56 | 0.94 |
|  |  |  |  | Bulb | 0* |  | 1 |
| **Transplanting date** | 123.3206 | 0.23 | 0.63 | Late October | 0.2804 | 0.59 | 1.32 |
|  |  |  |  | Early December | 0* |  | 1 |
| **Fungicide application** | 98.2558 | 25.06 | <.0001 | ≤ 4 times | -3.3253 | 0.84 | 0.035 |
|  |  |  |  | >4 times | 0* |  | 1 |

<sup>a</sup>Variables added into the model in order of presentation in table. <sup>b</sup>Unexplained variations after fitting the model. <sup>c</sup>LRT = Likelihood ratio test. DR = Deviance reduction. Pr = Probability of $\chi^2$ value exceeding the deviance reduction. $\chi^2$ = Chi-square. <sup>d</sup>Estimates from the model with all independent variables added. \* = Reference group. SE = Standard error

The independent variables that showed significant associations and some variables with higher deviance reduction were tested in a reduced multiple-variable model. The results of the analysis of deviation for each variable and variable class to incidence and severity, parameter estimates, standard error, and odds ratio are given in Tables 4 and 5. Onion downy mildew incidence and severity of >40% and >20% were highly associated with Fogera, and Libokemkem districts, land preparation (≤3 times), bulbing to maturity growth stage, onion growing after bulb, late October seedling transplanting and fungicide application (≤4 times spray) compared with their respective counterpart class variables (Tables 4 and 5). Among the tested independent variables, growth stage, land preparation, field size, seed source, previous crop fungicide application frequency were the most important variables in their association with onion downy mildew incidence and severity when entered first and last into the regression model respectively (Tables 4 and 5).

**Table 5.** Analysis of deviance, natural logarithms of odd ratio and standard error of onion downy mildew severity and likelihood ratio test on independent variables in northwestern Ethiopia, during the 2024/25 growing season.

| Variable<sup>a</sup> | Residual Deviance<sup>b</sup> | Downy mildew severity, LRT<sup>c</sup> |  | Variable class | Estimate<sup>d</sup> | SE | Odds ratio |
| --- | --- | --- | --- | --- | --- | --- | --- |
| | | DR | Pr > $\chi^2$ | | | | |
| **Intercept** | 180.1875 |  |  | Intercept | -1.1772 | 1.28 | 0.31 |
| **District** | 179.6648 | 0.52 | 0.2203 | Fogera | 4.1128 | 2.75 | 61.18 |
|  |  |  |  | Libokemkem | 2.4026 | 1.6 | 11.05 |
|  |  |  |  | Gondar Zuria | 1.4536 | 1.24 | 4.28 |
|  |  |  |  | Farta | 0* |  | 1 |
| **Growth stage** | 28.8016 | 18.85 | <.0001 | Vegetative | -2.8828 | 1.81 | 0.05 |
|  |  |  |  | Bulbing | -2.60 | 3.37 | 0.07 |
|  |  |  |  | Maturity | 0* |  | 1 |
| **Land preparation** | 95.3065 | 15.73 | <.0001 | ≤ 3 times | 4.5799 | 1.56 | 97.50 |
|  |  |  |  | > 3 times | 0* |  | 1 |
| **Field size (ha)** | 139.8549 | 6.55 | 0.0045 | ≤ 0.25 | -2.53 | 2.62 | 0.07 |
|  |  |  |  | > 0.25 | 0* |  | 1 |
| **Seed source** | 47.6509 | 25.12 | <.0001 | Farmer saved | -2.61 | 2.28 | 0.08 |
|  |  |  |  | Local market | 0* |  | 1 |
| **Plant population** | 54.3674 | 7.58 | 0.0001 | Low | -1.6370 | 2.82 | 0.19 |
|  |  |  |  | High | 0* |  | 1 |
| **Previous crop** | 96.9786 | 7.84 | <.0001 | Legume&cereal | -4.4270 | 1.57 | 0.011 |
|  |  |  |  | Bulb | 0* |  | 1 |
| **Fungicide application** | 70.0756 | 9.57 | 0.0020 | ≤ 4 times | -2.6799 | 0.95 | 0.07 |
|  |  |  |  | >4 times | 0* |  | 1 |

<sup>a</sup>Variables added into the model in order of presentation in table. <sup>b</sup>Unexplained variations after fitting the model. <sup>c</sup>LRT = Likelihood ratio test. DR = Deviance reduction. Pr = Probability of $\chi^2$ value exceeding the deviance reduction. $\chi^2$ = Chi-square. <sup>d</sup>Estimates from the model with all independent variables added. \* = Reference group. SE = Standard error. PSI = Percent severity index.

The probability of highest (>40%) onion downy mildew disease incidence was associated with Fogera and Libokemkem districts. The probability of the highest (>40%) disease incidence relation with Fogera and Libokemkem was about 3.25 and 2.79 times greater than the Farta district, respectively. Disease severity >20% showed high relationships with Fogera and Libokemkem compared to Farta district. The probability of highest (>20%) disease severity with Fogera and Libokemkem was about 61.18 and 11.05 times greater than Farta, respectively. maturity growth stage, less than three times plowing, fields covered with bulbs previously, seeds bought from local market, field size greater than 0.25ha showed a greater association with the highest disease incidence and severity.

Fungicide spray (≤ 4 times) and transplanting in late October showed a greater association with the highest incidence of the disease. In this regard, land preparation (≤ 3 times) was strongly associated with the highest disease incidence and severity about 6.29 and 97.5 times greater than more than three times plowing, accordingly. However, the lowest (≤ 40%) disease incidence and PSI (≤ 20%) exhibited a strong association with good land preparation, vegetative growth stage, farmer saved seeds, less than 0.25ha field size, cereal previous crop (Tables 4 and 5). Transplanting early December and greater than four times fungicide spraying were less associated with only (≤ 40%) disease incidence (Table 5).

## 4. DISCUSSION

Onion downy mildew is caused by the fungus *Peronospora destructor* causing serious epidemics in Ethiopia’s humid and temperate onion-growing regions. Regardless of environmental and onion management practices, onion downy mildew was prevalent in all surveyed fields. In northwestern Ethiopia, yield loss due to downy mildew has been recorded up to 80% of the total production. Due to favorable environmental conditions, downy mildew was more abundant and widespread in northwestern districts of Ethiopia [27, 19]. Onion downy mildew disease incidence and severity varied across agro-ecological conditions. The highest disease intensity was assessed from the Fogera district than the other districts. The initial inoculum, cultural practice, and environmental factors may be the cause of the variance in disease epidemics among the surveyed districts. This suggests that biological, cultural, and environmental factors had a major impact on the development and spread of onion downy mildew disease. Factors including as inoculum load, farming practices and environmental variables might also affect the disease levels in different each districts. Because of several agro-ecological conditions, the main and irrigation onion production systems at Fogera Ethiopia showed the maximum onion downy mildew disease severity [41].

Onion downy mildew is naturally more common at higher elevations, especially in regions with cool temperatures and moisture but the disease intensity were higher in fields at altitudes ≤ 1800m.a.s.l. at the Fogera and Libokemkem districts respectively than onion fields assessed at altitudes above 800m.a.s.l. Because, these areas during onion growing season had optimum temperature to pathogen, high night, and morning humidity and collaborating with other agro ecological variables (figure 2). These combined conditions were hot spot for pathogen infection leads to disease development and epidemics. The pathogen sporulated during night, with sporangia maturing in the early morning and being released throughout the day. The sporangia were favored by a wide range of temperatures and a lot of moisture [42, 43, 44]. Favorable early morning temperatures and the few hours of high humidity have been attributed for a major mildew outbreak in rainless weather [45, 46].

Obviously, the effects of onion downy mildew might vary depending on the stage of growth, from vegetative to maturity. In this study, onion downy mildew disease epidemic was higher at bulbing to maturity stages than the vegetative stage and associated with higher disease incidence and severity. The presence of older leaves, a decrease in the start of new leaves, and a loss of crop resistance to the disease make onions more susceptible to systemic infection at maturity. The bulbing to maturity stage of onion plants was shown to be particularly susceptible with older leaves present because new leaf initiation is stopped and crop resistance to the disease is lost have been reported[47,19].

Regarding to the previous crop, Onion downy mildew outbreaks were significantly associated with the planting of onions following bulbs. The occurrence and severity of onion downy mildew may be significantly influenced by past crop history. Spores that overwinter in crop debris can serve as a source of inoculum for just emerging plantings. Seasonal outbreaks of onion downy mildew may be significantly influenced by the survival of overwintering structures. It is known that *Peronospora destructor* naturally develops oospores that can persist in the soil for a long time [48,49]. The initial inoculums may distributed in to field through irrigation, wind, plowing and by another mechanisms and increasing the risk of infection in subsequent crops. This could lead to an increase in epidemics and the occurrence of onion infection by the disease. The main source of inoculum for subsequent bulb crops is diseased debris that contains pathogen fruiting bodies in the field of crops and onion seeds [50, 51, 52].

Fields ploughed more than three times related with less disease intensity. Frequent ploughing may disperse the organic matter in the soil that is appropriate for the robust growth of crops, making them less vulnerable to pathogens. It is also used to expose the main inoculum to solar energy, which breaks the initial inoculum population in the soil. This may result in the decrement of disease epidemics. Previous study by [53], reported that many times plowing reduce relatively disease epidemics than with few times ploughed plots. During the current field survey, Fields with a high population density had the highest mean disease intensity. This may mainly be related to a microenvironment that promotes disease outbreaks due to higher humidity, decreased air circulation, and high plant nutrient completeness. Previously scholars [54] reported that the development and outbreak of plant diseases are influenced by the morphology, canopy, and density of plants.

Regarding field size, onion growing in small areas (≤ 0.25ha) had less disease intensity than large cultivated area (*>*0.25ha). The association of higher disease epidemic in large field size might be create a more favorable environment for downy mildew because the fields are more humid, have less air circulation, and are more susceptible to sporangia spreading from one area of infection to another due to several factors, particularly through irrigation and wind. Due to it’s easily dissemination, the outbreak of diseases was abundant in large onion fields [55,56] showed how easily downy mildew sporangia may be distributed and moved from one field to another through wind. High disease incidence and severity were observed with onion fields transplanting in late October, particularly when combined with other factors. This is a result of having an environment that is conducive to the establishment of pathogens and the development of disease. During this season, these studied sites had low temperatures and intermittent rainfall, which may have played a significant role in the development of disease. The periods of rainy season, high moisture and humidity promotes the disease development [57, 58, 59].

Fungicide spraying (≤ 4 times) on onion fields resulted in greater disease development compared with fungicide application (> 4 times), The application that occurs the least frequently Fungicides might not disrupt the life cycle of the pathogen and spread unevenly in order to prevent the development of diseases and this may lead pathogen adaptation against chemicals resulted in diseases outbreak. Therefore, frequent application Fungicides are often used to prevent the germination, growth, or multiplication of the pathogen, which has suppressed the spread of onion downy mildew. High spraying frequency is the primary factor that determines the success of chemical control [60]. Plots with frequent scheduled fungicide application to ensure good yields and high crop quality showed the lowest average disease severity and highest yield [61, 62]. The present study shown that downy mildew damages bulb crops in areas with high humidity and moisture content; as a result, the disease was more affected in the Fogera district.

## 5. CONCLUSION

Onion downy mildew disease was distributed variably in each of the districts that were surveyed. The disease was widely distributed and prevalent in all surveyed areas with high incidence and severity in the Fogera, followed by Libokemkem district. The independent variables, such as transplanting of onion until late October, maturity growth stages, poor land preparation, seeds bought from market, large field size, more than four times fungicide applications and bulb previous crop were shown to be crucial variables that contributed to the downy mildew disease outbreak and exhibited a significant association with the highest diseases intensity.

The multiple-reduced regression model analysis identified significant associations between district, crop growth stage, land preparation, plant population, field size and previous crop history, and downy mildew incidence and severity. Additionally, the model assessed the relative relevance of the biophysical parameters under study, which may indicate that some variables, either alone or in combination, contributed to the development of the downy mildew epidemic in the study areas. In this regard, among the highly onion producing areas, Because of downy mildew diseases, the areas of Fogera and Libokemkem were the most problematic for growing onions. The study indicated that transplanting on early December, growing of onion in previously cereal cultivated land, good land preparation, and 4-6 times frequent fungicide applications may be regarded as useful options for management to mitigate the disease’s effect on onion crops.

## 7. ETHICAL STATEMENT

This material is the authors own original work, which has not been previously published elsewhere and is not currently being considered.

## 8. DATA AVAILABILITY STATEMENT

The datasets generated and analyzed during the current study are available from the corresponding author on reasonable request.

## 10. AUTHOR CONTRIBUTIONS

Mitku Bitew: Conceptualization, investigation, formal analysis, writing-original draft and writing-review.

Addisu Mandafro: Conceptualization; supervision; validation; writing-review and editing. Yehizbalem Azmeraw: Conceptualization; supervision; validation; writing-review and editing.

